# Rethinking the importance of the structure of ecological networks under an environment-dependent framework

**DOI:** 10.1101/219691

**Authors:** Simone Cenci, Chuliang Song, Serguei Saavedra

## Abstract

A major quest in network and community ecology has been centered on understanding the importance of structural patterns in species interaction networks—the synthesis of who interacts with whom in a given location and time. In the past decades, much effort has been devoted to infer the importance of a particular structure by its capacity to tolerate an external perturbation on its structure or dynamics. Here we demonstrate that such a perspective leads to inconsistent conclusions. That is, the importance of a network structure changes as a function of the external perturbations acting on a community at any given point in time. Thus, we discuss a research agenda to investigate the relative importance of the structure of ecological networks under an environment-dependent framework. We hypothesize that only by studying systematically the link between network structure and community dynamics under an environment-dependent framework, we can uncover the limits at which communities can tolerate environmental changes.

“*Variation stands out as the only meaningful reality*”

Stephen J. Gould.

## Introduction

Since the beginnings of modern network theory [36, 37], studies have assessed the importance of particular network structures (e.g., exponential or scale-free networks) by their capacity to tolerate an external perturbation acting on their structure or dynamics (e.g., a random or targeted sequential removal of nodes, Albert et al. 1). This has paved the way for a similar research agenda in network and community ecology [3, 39]. In particular, theoretical studies have been investigating this importance by quantifying the effects of network structure on community persistence [34]. To capture these effects, a typical approach has been centered on randomly removing species (removing interactions or sampling randomly model parameters) and comparing the extent to which different network structures avoid additional species extinctions. This tolerance has then been taken as evidence for a structure’s advantage, disadvantage, or lack of any importance over other structural patterns [26, 47]. However, the large number of degrees of freedom involved in these analyses (e.g., parameter values, choice of perturbation) has been a central limitation. In fact, it is unclear the extent to which such conclusions can be generalized [21, 45]. Therefore, the question has become whether it is possible at all to infer the importance of a network structure through its capacity to tolerate external perturbations [42].

In this manuscript, we use a simple example to demonstrate that the tolerance to external perturbations of different network structures under the same dynamics can quickly change as a function of the type, direction, and magnitude of the perturbations. That is, the importance of a network structure depends on the external perturbations faced by a community at any given point in time [8, 14, 52]. Thus, if studies focus on a specific set of external perturbations in order to infer the general importance of a network structure, it would lead to inconsistent conclusions. This implies that the importance of a given network structure in a community should always be understood in relation to local environmental settings. In this line, we propose and discuss a research agenda to investigate the relative importance of the structure of ecological networks under an environment-dependent framework. We strongly believe that this new synthesis can move the field of ecology towards a more systematic and predictive science [40].

## Linking network structure and persistence

To investigate the tolerance of an ecological network to external perturbations, studies have been linking the structure of interaction networks with community persistence. Traditionally, this structure has been derived from the topology of species interaction networks, i.e., the binary representation of who interacts with whom in a given location and time [3, 39]. This topology can be represented by a binary matrix, whose elements denote the presence or absence of a direct interaction between two species. In order to talk about the structure of a network, it has also been necessary to talk about the lack of structure in a network [37]. In this context, random networks (or null models) have been the gold-standard benchmark of no structure. Generally, these random networks are simple ensembles of binary matrices with a given number of 1s and 0s randomly shuffled. Statistically significant deviations from these random networks have been taken as a sign of a structure in an observed interaction network [60]. Note that the characterization of a structure is not restricted to its topology, and many different definitions (e.g., weighted instead of binary patterns) and null models can be used [51]. Yet, the conceptual framework is exactly the same.

This framework has revealed that many realworld networks exhibit a distribution of interactions that depart from null expectations [37]. For example, in network ecology, two of the structures that have captured most of the attention are modular and nested structures [3, 39]. Modular structures are those in which groups of species have many interactions among them, but few interactions with the rest of the species in the network. Nested structures are those where highly connected species interact with both highly connected and poorly connected species, while poorly connected species interact almost exclusively with highly connected species. Thus, many studies have been interested in understanding the existence and importance of such network structures through their potential links with community persistence.

Formally, community persistence corresponds to the capacity of a given community to sustain positive abundances for all its constituent species [22]. In many cases, this definition has been relaxed and taken as the fraction of species that can sustain positive abundances in a community when subject to some initial conditions or external perturbations [7, 25, 39]. In the absence of empirical data, the challenge has been how to model the temporal evolution of species abundances and their response to external perturbations [42]. Therefore, some studies have analyzed community persistence by assuming a random or targeted sequential removal of species (or interactions) in an interaction network, and considered extinctions (i.e., when abundances go to zero) whenever a species is left without interactions [39]. Other studies have used population dynamics models to investigate the fraction of species that ends up with positive abundances at equilibrium under some random initial conditions [42]. Similarly, using population dynamics models, other studies have systematically investigated the range of parameter values (initial conditions) compatible with positive abundances of all species in a community [44]. Overall, regardless of the method employed, community persistence has been broadly defined as the capacity of a community to avoid species extinctions.

One can now link the two concepts above and ask, for example, to what extent modular or nested structures can increase the capacity of communities to avoid species extinctions under random perturbations. In fact, this has been a recurrent question in network and community ecology [3, 39, 47]. To address this question, studies have fixed a community size, changed the structure systematically by re-arranging the interactions of the networks, adopted a measure of community persistence, introduced some external perturbation or condition, and investigated how this measure of community persistence changes as a function of a network structure [4, 26, 42, 54, 57]. Importantly, these theoretical studies have shown significant associations between network structures and community persistence under given external perturbations.

However, the reading of these findings has many times led to a belief that a structure is either advantageous (important) or not for a community [47]. For example, modular and nested structures have been positively associated with community persistence in antagonistic and mutualistic communities, respectively [57]. Thus, changes of a presumed important network structure in a community across temporal or environmental gradients have been directly translated to changes in robustness [16, 56, 64]. Moreover, this view has led other studies to suggest that the absence of a given structural pattern in a community can be related to the lack of importance of such network structure overall [26, 55]. Yet, all these generalizations are derived from particular scenarios of external perturbations, and currently it is unknown whether these results are consistent under a more systematic analysis [21, 45].

## Inconsistent conclusions about the importance of network structure

To illustrate how naive simulations of external perturbations can lead to inconsistent conclusions about links between network structure and community persistence we follow a structural stability approach [42, 58]. This approach is particularly useful for our purposes as it allows us to focus on how the qualitative behavior of a dynamical system changes as a function of the parameters of the system itself. For example, the dynamics of the system can be approximated by a population dynamics model [10]. Then, the qualitative behavior of this dynamical system can be translated into a given measure of community persistence. Thus, one can investigate the extent to which different interaction networks can tolerate external perturbations (changes in parameter values) without pushing species towards extinction.

This approach can also be graphically represented. For example, for a 2-species community, the axes of 1 represent the 2-dimensional parameter space of species vital rates. The colored regions correspond to the set of those vital rates compatible with positive species abundances (the necessary condition for community persistence). The size and shape of this region depend upon network structure [11]. This region is typically called the feasibility domain of a community [32, 42]. The open and colored symbols represent some initial and final parameter values (after a hypothetical external perturbation), respectively. Rows correspond to the same external perturbation under two different network structures (the reader can think of any type of structures). Columns correspond to the same network structure under two different external perturbations. Note that positive species abundances will be satisfied as long as the parameter values fall inside the feasibility domain. If we were to focus on the first row only, we would conclude that structure 1 is more robust that structure 2. However, if we were to focus on the second row, then we would conclude the opposite. Similarly, if we were to focus on each column separately, we would arrive to contrasting conclusions. Moreover, these inconsistent conclusions can be repeated by moving the perturbation (parameter values) to almost any other direction. That is, there is no conceptual support to think of either a positive or negative association exclusively, especially not by focusing on a single type of perturbation.

More systematically, let us take the classic Lotka-Volterra (LV) dynamics 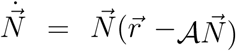 as a toy model. These dynamics are useful as one can directly associate the structure of the feasibility domain with the network structure [46]. However, note that results of our discussion extend to a larger class of population dynamics models with nonlinear functional responses [12]. In the LV model, the abundances of species are represented by the *n*-dimensional vector 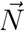, where *n* corresponds to the community size. The temporal evolution of species abundances 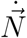 is a function of the abundances at any given point in time 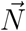, the vector of intrinsic growth rates (vital rates) of species 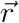, and the interaction matrix 𝒜 [10]. Note that the interaction matrix is a quantitative description of the interaction network, while the values of intrinsic growth rates are inherently linked to environmental conditions [8, 14, 31]. If we take our measure of community persistence as that in which there are no extinctions at equilibrium (i.e., 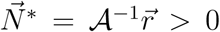), then we can see that this condition will be satisfied as long as the vector of intrinsic growth rates 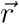 falls inside a feasibility domain constrained by the interaction matrix *A* [42, 44]. Formally, this domain is defined by

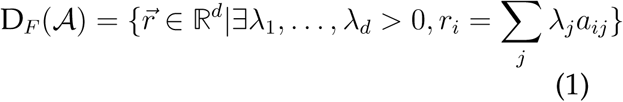

To quantitatively illustrate the inconsistent conclusions about the importance of the structure of interaction networks through their tolerance to external perturbations, we test the association of community persistence with modular and nested structures under LV dynamics. Yet, we need to stress that our approach can be applied to any combination of structures, perturbations, and models. We measure community persistence as the capacity of a particular structure to avoid extinctions. We build interaction networks on communities of 21 species (this number allows us to easily divide the network into modules, but different dimensions generate the same qualitative results). Interactions are distributed among the species so that there is a clear distinction between the two types of structures analyzed (see 2 for a graphical representation). For comparison purposes, the elements of the interaction matrix 𝒜 are taken from a normal distribution with parameters chosen such that the resulting interaction matrices for each structure have same mean and standard deviation. In the absence of an interaction between two species, the corresponding entry in the interaction matrix is zero. Communities (each with a different type of structure) are initialized inside the feasibility region by fixing a lognormal distribution of species abundances 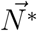 [5], and then finding the corresponding vectors of intrinsic growth rates i.e., 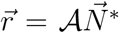 [43]. Once the communities (with the different structures) are initialized with all species present 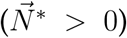, we introduce random and directional perturbations on either the interaction matrix or the vector of intrinsic growth rates. While random perturbations act on all the elements of the interaction matrix or the vector of intrinsic growth rates, directional perturbations act on one single column or element. These changes are equivalent to random and targeted perturbations either on the interactions or nodes of a network [45, 46]. After the perturbations, we compute the new equilibrium solution with the changed parameters. Then, we record which community (network structure) avoids extinctions. We repeat this process 5000 times.

Figure 2 shows the estimated community persistence (number of times a given structure avoids extinctions) derived for each combination of structure and perturbation. The first row corresponds to the community persistence under random perturbations acting on either the interaction matrix (Panel A) or the vector of intrinsic growth rates (Panel B).

The second row corresponds to the community persistence under directional perturbations acting on either the interaction matrix or on the vector of intrinsic growth rates of the most and least connected species. In the same line as in Figure 1, if we were to focus on the first row only, we would conclude that modular and nested structures are more robust under perturbations acting on the interaction matrix and intrinsic growth rates, respectively. However, if we were to focus on the second row, then we would conclude the opposite (Panels C and F), or simply that there is no difference between the structures (Panels D and E). Similarly, if we were to focus on each network structure separately, we would arrive to contrasting conclusions as a function of the perturbations. Note that these inconsistencies are not exclusive to the perturbations here analyzed. Overall, these simple conceptual and quantitative analyses demonstrate that the association of a given network structure with community persistence completely depends on the type, direction, and magnitude of perturbations.

**Figure 1:**
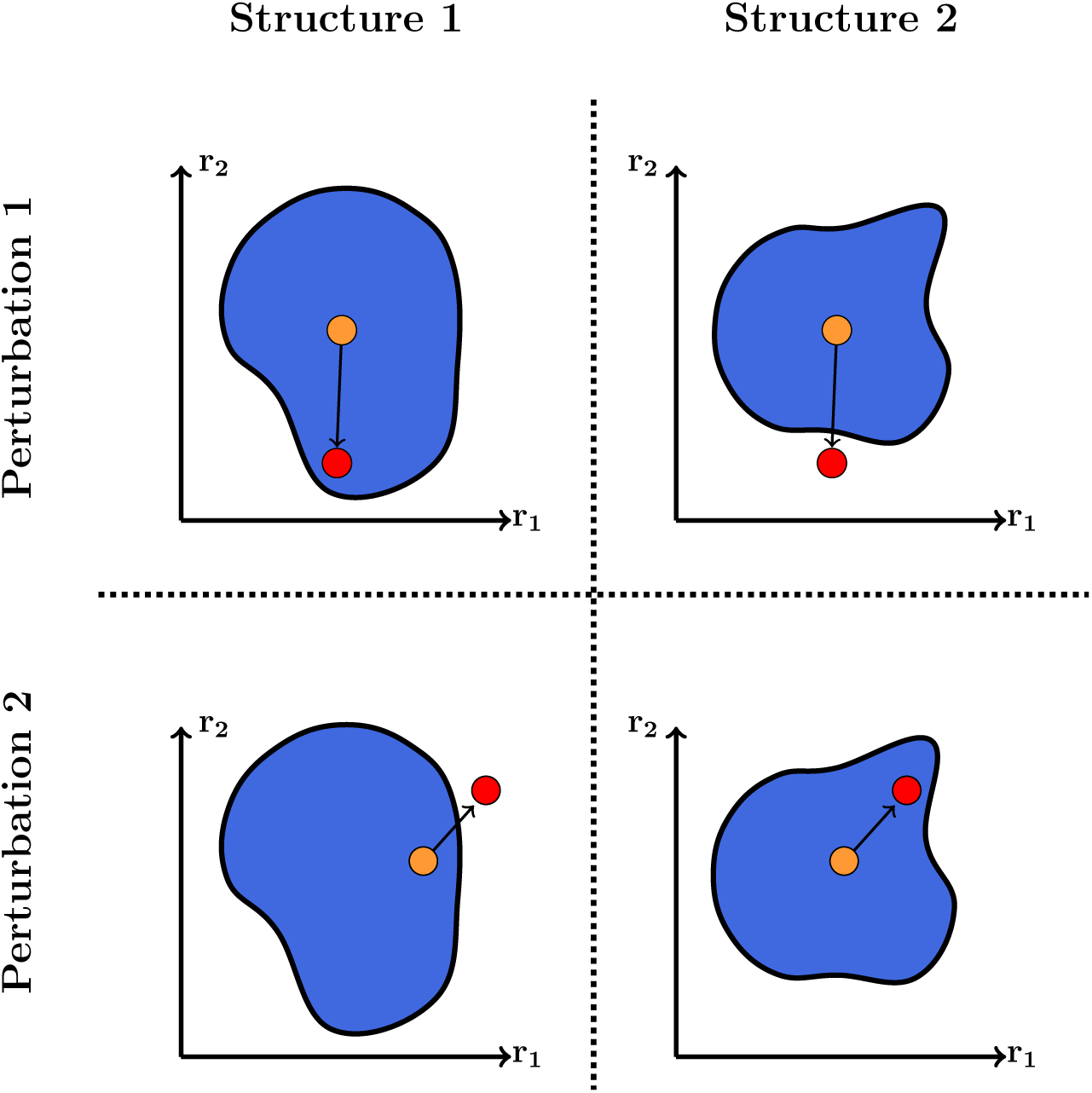
Linking external perturbations, network structures, and community persistence. The blue region represents the feasibility domain (parameter space compatible with community persistence) of a population dynamics model. The orange and red circles represent a vector of species intrinsic growth rates 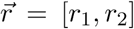 before and after a hypothetical perturbation, respectively. The necessary condition for community persistence is to have a vector of intrinsic growth rates within the feasibility domain (as we show on the top-left and bottom-right panels). The cartoon shows that not only the structure of an interaction network is important for community persistence, but also the direction of the perturbation. In fact, just by changing the direction of the perturbation, one may not observe community persistence under the same network structures (as we show on the top-right and bottom-left panels). That is, structure *per se* says little about community persistence if not seen in the light of its local environment.

**Figure 2:**
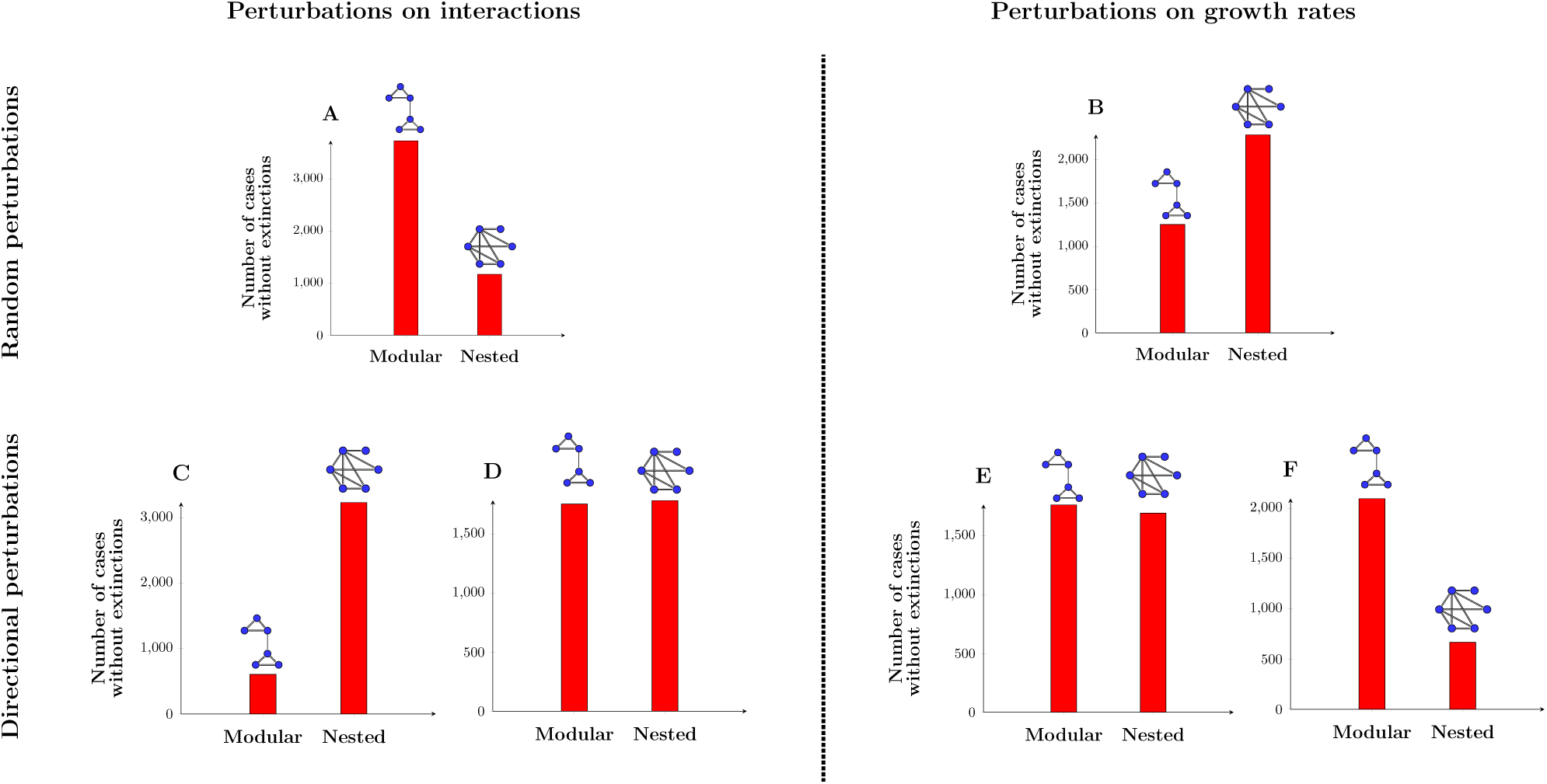
Inconsistent conclusions about the importance of network structures. As an example, we show the response of two different network structures (a modular and a nested structure) to different external perturbations (see text for details on the simulations performed using a Lotka-Volterra competition model). All communities are initialized inside the feasibility domain having the same species abundance distribution (see text for details). The y-axis corresponds to the number of times (out of 5000) that the community tolerated a perturbation (i.e., no species goes extinct). That is, the large the bar, the more tolerant the network structure. Panel **A** corresponds to random perturbations on species interactions. Panel **B** corresponds to random perturbations on species intrinsic growth rates. Panels **C** and **D** correspond to directional perturbations on species interactions. That is, only the values of one column of the interaction matrix are changed in each case. Finally, Panels **E** and **F** correspond to directional perturbations on species intrinsic growth rates. That is, only one growth rate of one species is changed in each case.

## A plea for an environment-dependent framework

As it is known, species interaction networks are the result of different evolutionary and ecological processes acting at the individual and the collective level [59]. Because these processes typically yield to adaptation to local environments [20, 61], it becomes useful to think about the relative importance of a network structure under a particular environmental setting. That is, the importance of a network structure should be studied under an environment-dependent framework [13, 52]. This research agenda can be achieved by using environment-dependent parameters as the link between community dynamics and environmental conditions [8, 43]. The goal will be to investigate the range of environmental conditions compatible with community persistence under a given network structure, the expected environmental conditions in a given location, and their overlap. Below, we explain this environment-dependent framework in more detail.

The first step is to systematically study the set of environmental conditions tolerated by a community with a given network structure. This requires to link community dynamics and environmental conditions, which can be achieved through specifying environment-dependent parameters in a model (e.g., species carrying capacities) [8, 14]. That is, instead of studying the tolerance of a network structure as a function of random environmental conditions, we propose to study the set of conditions (range of environment-dependent parameter values) compatible with the persistence of the community. This set is what we have previously called the feasibility domain. Recall that this domain is defined by the particular dynamics and network structure of a community [42]. As an illustration, Figure 3A shows the feasibility domain of two different network structures under the same population dynamics. Importantly, in many cases, the size and shape of the feasibility domain can be analytically investigated [44].

**Figure 3:**
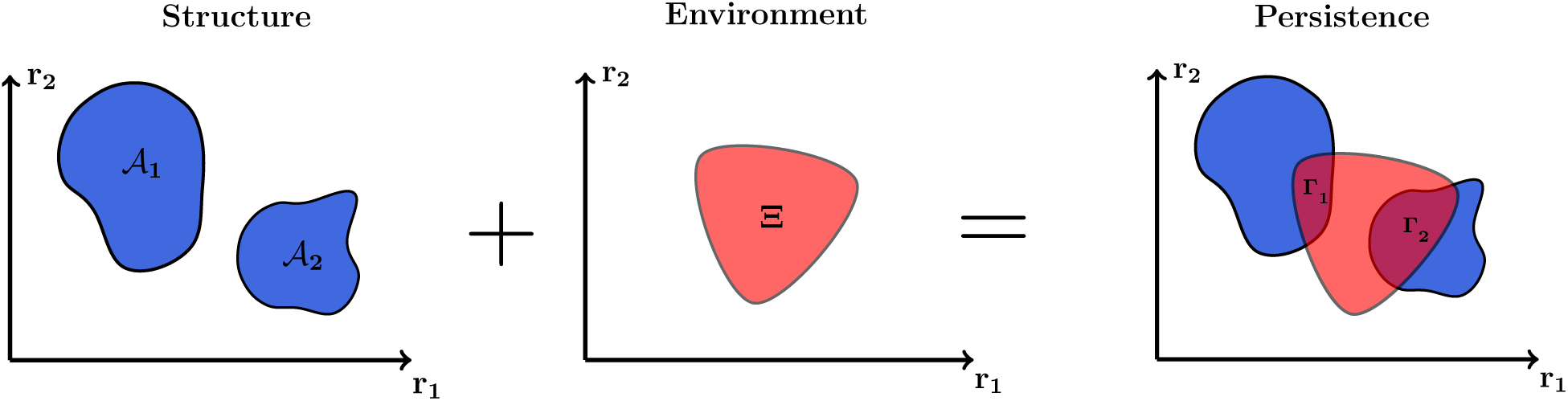
The relative importance of network structures under an environment-dependent framework. This cartoon illustrates the extent to which the importance of network structures depends on local environmental settings. The left panel shows two different feasibility domains (range of environment-dependent parameter values compatible with community persistence) of a hypothetical 2-dimensional system. These domains are generated by two different network structures characterized by their corresponding interaction matrices. Note that the left domain is larger than the one on the right. The middle panel corresponds to a hypothetical characterization of an environment in a given location (e.g., set of environmental conditions over a period of time) as a function of environment-dependent parameters. This region of parameters can also be represented by a distribution. The right panel corresponds to the overlap between the environment and the two network structures (i.e., community persistence). Note that the smaller feasibility domain 𝒜 _2_ can have a larger overlap with the environment Ξ (i.e., Γ_2_ *>* Γ_1_ in the Figure), providing a higher probability of persistence for the community.

The second step involves estimating the range of environment-dependent parameter values compatible with the environment in a given location (i.e., the local environment over a period of time) [14, 40]. For example, it may be possible to map the influence of environmental conditions; such as temperature, humidity, or soil composition, into species carrying capacities [17, 19, 28, 30, 35, 52]. Similarly, species carrying capacities can be linked to the total amount of available resources per consumption rate, where monocultures have been used as an experimental ground to estimating such values [10]. Note that this estimation could also be represented by a probability distribution for the carrying capacities [8]. That is, a measure of which type of carrying capacity are more likely to be observed in a given environment. As an illustration, Figure 3B shows, a hypothetical environment in a given location as a function of environment-dependent parameters. Note that this characterization would also require knowledge about climatic conditions or the use of weather models [49], as well as knowledge about how individual species respond to those changes. While this task could be challenging, both new theoretical [2, 6, 15, 23, 24, 41] and empirical studies [9, 50, 62] are providing a good guideline towards this goal. Overall, the task is to characterize a representative set of potential environmental conditions for a given location rather than an arbitrary set of random external perturbations.

The third step corresponds to merging steps one and two. Because it is virtually impossible to know the type, direction, and magnitude of environmental conditions acting on a community at every given point in time; it becomes useful to study the extent to which the environmental conditions compatible with a given network structure overlap with the environment faced by a community in a given location. This overlap corresponds to the proportion of the feasibility domain of a community that is inside the set of environment-dependent parameter values expected in a given location (for a graphical example see Figure 3C). Formally, this proportion can be defined by the ratio of the following volumes: Γ(*D*_*F*_ (𝒜_*i*_) ∩ Ξ_*j*_) = vol(*D*_*F*_ (𝒜_*i*_) ∩ Ξ_*j*_)*/* vol(Ξ_*j*_), where *D*_*F*_ (𝒜_*i*_) corresponds to the feasibility domain of a community *i* (network structure) and Ξ_*j*_ corresponds to the distribution *j* of environment-dependent parameter values expected in an environment. In other words, community persistence should always be measured under an environment-dependent framework [31]. This definition implies two important concepts: (1) Under the same environment Ξ_*j*_, differences among structures (𝒜 _1_ and 𝒜 _2_) can reflect differences among feasibility domains, and, in turn, differences among community persistence (Γ_*ij*_). (2) Under different environments (Ξ_1_ and Ξ_2_), differences among feasibility domains do not directly reflect differences between community persistence. That is, a community with a small feasibility domain can have a much greater community persistence that a community with a large feasibility domain if the environment happens to overlap more with the domain of the small one (see 3C for an illustration). While this approach certainly does not remove the existence of inconsistent conclusions under different environments (e.g., moving from Ξ_1_ to Ξ_2_), it allows us to better characterize the link between network structures and environmental settings.

## Final thoughts

We have shown that it is necessary to rethink the importance of the structure of ecological networks under an environment-dependent framework; otherwise, we may miss the forest for the trees. Because local adaptation is the *leitmotiv* of natural selection, a plausible hypothesis is that network structures are emergent responses of communities to their environment [11, 14, 29, 31, 33, 38, 61]. That is, there is no one better network structure than other in general. However, there can be one network structure more tolerant than other structure to a particular environment. This also implies that a change in network structure is neither advantageous nor detrimental *per se*. Changes in network structure can be the result of different factors, such as: external perturbations, ecological or evolutionary processes, invasions, and extinctions, among others. Similarly, these changes can be adaptive or non-adaptive, revealing that changes of a presumed important network structure in a community across temporal or environmental gradients cannot be directly translated to changes in robustness. In fact, studies have already shown contrasting effects of structural changes on adaptability [18, 27]. Thus, again, these analyses can only make sense within an environmental context. Therefore, the lack of a structural pattern in one or several communities does not imply that such structure has no importance at all. It may only imply that such structure is not particularly advantageous under the current environmental settings.

While we have developed a non-exhaustive quantitative exercise of the many possible structures, dynamics, and perturbations, we have shown clear counter-examples of how conclusions about the importance of a network structure derived from particular combinations are inconsistent. Thus, because it is virtually impossible to know *a priori* all the characteristics of future perturbations [48, 53, 63]; we need to abandon a dichotomous view, and systematically link network structure and community persistence under an environment-dependent context. While the approach we have outlined in this work is by no means the only possible one, we hope it can be used as a guideline towards a better understanding of both the existence of structural patterns in ecological networks and how communities may respond under future scenarios of climate change.

